# Theta-modulation drives the emergence of network-wide connectivity patterns underlying replay in a model of hippocampal place cells

**DOI:** 10.1101/118489

**Authors:** Panagiota Theodoni, Bernat Rovira, Yingxue Wang, Alex Roxin

**Affiliations:** Computational Neuroscience Group, Centre de Recerca Matemàtica, Campus de Bellaterra, Edifici C, 08193 Bellaterra, Spain; NewYork University Shanghai, 1555 Century Ave, Pudong, Shanghai 200122, China; NYU-ECNU Institute of Brain and Cognitive Science at NYU Shanghai, 3663 Zhongshan Road North, Shanghai 200062, China; Max Planck Florida Institute for Neuroscience, 1 Max Planck Way, Jupiter, Florida 33458, USA

## Abstract

Place cells of the rodent hippocampus fire action potentials when the animal traverses a particular spatial location in a given environment. Therefore, for any given trajectory one will observe a repeatable sequence of place cell activations as the animal explores. Interestingly, when the animal is quiescent or sleeping, one can observe similar sequences of activation, although at a highly compressed rate, known as “replays”. It is hypothesized that this replay underlies the process of memory consolidation whereby memories are “transferred” from hippocampus to cortex. However, it remains unclear how the memory of a particular environment is actually encoded in the place cell activity and what the mechanism for replay is. Here we study how plasticity during spatial exploration shapes the patterns of synaptic connectivity in model networks of place cells. Specifically, we show how plasticity leads to the emergence of patterns of activity which represent the spatial environment learned. These states become spontaneously active when the animal is quiescent, reproducing the phenomenology of replays. Interestingly, replay emerges most rapidly when place cell activity is modulated by an ongoing oscillation. The optimal oscillation frequency can be calculated analytically, is directly related to the plasticity rule, and for experimentally determined values of the plasticity window in rodent slices gives values in the theta range. A major prediction of this model is that the pairwise correlation of place cells which encode for neighboring locations should increase during initial exploration, leading up to the critical transition. We find such an increase in a population of simultaneously recorded CA1 pyramidal cells from a rat exploring a novel track. Furthermore, in a rat in which hippocampal theta is reduced through inactivation of the medial septum we find no such increase. Our model is the first to show how theta-modulation can speed up learning by facilitating the emergence of environment-specific network-wide patterns of synaptic connectivity in hippocampal circuits.

## Introduction

As an animal explores in any given environment, place cells in the hippocampus fire selectively at particular locations^1–3^, known as the “place-fields” of the cells. Furthermore, the place fields of an ensemble of place cells remap to completely new positions with respect to one another if the animal enters a distinct environment^4–6^. Sequential place cell activation during exploration therefore acts as a unique fingerprint for each environment, providing information needed for navigation and spatial learning. The spontaneous replay of such sequential activation, which occurs within sharp-wave/ripples (SWRs) during quiet wakefulness^7–9^ and sleep^10,11^, suggests that the animal has formed an internal representation of the corresponding environment, presumably during exploration^12^. Synaptic plasticity is the clear candidate mechanism for this formation. Nonetheless it remains unclear how changes in the synaptic connectivity are coordinated at the network level in order to generate well-ordered sequences spontaneously.

An additional prominent physiological signature of exploratory behavior in the hippocampus is the theta rhythm (4-12Hz). Decreases in theta power due to lesions of the medial septum strongly reduce performance in spatial-memory based tasks^13–15^ and other hippocampal dependent tasks^16,17^, although they do not eliminate place fields on linear tracks^18,19^. Despite this, lesioned animals can still reach fixed performance criteria given enough time^13,16^. This shows not only that theta is important for learning, but also suggests a potential mechanism. Namely, it may act as a dial for the learning rate, presumably by modulating the network-wide coordination of synaptic plasticity. How this occurs is unknown.

Here, we develop a model that explains how synaptic plasticity shapes the patterns of recurrent connectivity in a hippocampal circuit as an animal explores a novel environment. We show that temporally asymmetric plasticity rule^20,21^ during motion-driven sequential place-cell activity leads to the formation of network structure capable of supporting spontaneous bursts. These bursts occur in the absence of place-field input, i.e. during awake-quiescence or sleep. Importantly, the spatio-temporal structure of the bursts undergoes a sharp transition during exploration, exhibiting well-ordered replay only after a critical time of exploration in the novel environment. The underlying rate of this plasticity process, and hence the critical time, is strongly modulated through external oscillatory drive: for very low and high frequency the rate is near zero, while for an intermediate range, set by the time-scale of the plasticity rule, it is higher by several orders of magnitude, allowing for learning on realistic time-scales. Our theoretical findings lend support to the hypothesis that the theta rhythm accelerates learning by modulating place-cell activity on a time-scale commensurate with the window for plasticity. This maximizes the growth rate of network-wide patterns of synaptic connectivity which drive spontaneous replay. Finally, we confirm a main prediction from the model using simultaneous recordings of hippocampal place cells from a rat exploring a novel track. Namely, pairwise correlations between cells with neighboring place fields show a sharp increase over the span of several minutes at the outset of exploration. Furthermore, in a rat in which the medial septum is partially inactivated via muscimol injection, which strongly reduces theta modulation, this increase is not seen.

## Results

### Plasticity during exploration of a novel environment leads to a transition in the structure of sharp-wave events

We modeled the dynamics of hippocampal place cells as an animal sequentially explored a series of distinct novel ring-like tracks, Fig.1a. The model consisted of recurrently coupled, excitatory stochastic firing rate neurons, each of which received a place-specific external input on any given track, and inhibition was modeled as a global inhibitory feedback, see *methods* for model details. To model the global remapping of place fields from one track to another, we randomized the position of the peak input, i.e. the place field, of each neuron. In this way, the ordering of cells according to their place field location on one track was random and uncorrelated with their ordering on any other track, see Fig.1b.

**Figure 1.**
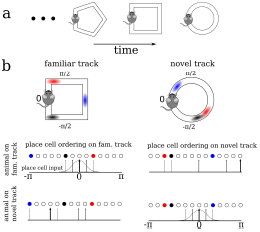
A schematic description of the model. (a) We model the sequential exploration of a number of distinct linear tracks. (b) The network consists of N place cells. The ordering of the place fields is randomly remapped on each track. Therefore, if the cells are properly ordered in any given environment the place field input is represented by a spatially localized bump of activity (upper left and lower right). Sequential activity on a familiar track would look random given an ordering on a novel track, and vice versa (upper right and lower left respectively).

We were interested in knowing how the exploration of a novel environment affected the pattern of synaptic connectivity between place cells via a temporally asymmetric plasticity rule, and how this in turn shaped the activity. While the plasticity rule we use is formally referred to as “spike-timing dependent” (STDP)^22–24^, our model neurons generate spikes stochastically as Poisson processes. Therefore the exact spike timing itself does not play any role but rather only the time variations in the underlying firing rate. Recent theoretical work has shown that the plasticity induced by a number of plasticity rules, including heuristic STDP rules, is in fact dominated by such firing rate variations when irregular, in-vivo like activity is considered^25^. We also considered a “balanced” plasticity rule, for which the degree of potentiation and depression were the same, on average, given constant firing rates. Such a rule is a means of avoiding fully potentiating or fully depressing all synapses on long time-scales, i.e. it allows for the generation and maintenance of structured synaptic connectivity. Alternatively, we could have considered an unbalanced plasticity rule with additional stabilizing mechanisms^26^.

In order to study “typical” place-cell dynamics during exploration we first exposed the network to a series of distinct tracks until the matrix of recurrent synaptic weights no longer depended on its initial state. When the trajectory and velocity of the virtual animal was stochastic and distinct on different tracks, the resulting evolution of the synaptic weights also exhibited some variability from track to track, see Fig.S1. In simulations with constant velocity this variability largely vanished, see Fig.S6a. After exploring 10 tracks, we exposed the network to another novel track and studied the dynamics in detail; because the exploration process had already become largely stereotyped (despite some variability), the dynamics on the novel track reflected what would be seen qualitatively during the exploration of any other novel track in the future, i.e. it was dependent only on the learning process itself and not the initial state of the synaptic weight matrix.

In our simulations, we modeled the movement of a virtual animal around the track by varying the external inputs to place cells. Specifically, the external input was maximal for cells with place fields at the animal’s current position and minimal for cells with place fields at the opposite side of the track, with the input decaying smoothly for intermediate locations like a cosine curve, see *methods* for details. We modeled the motion of the animal by taking the velocity to be an Ornstein-Uhlenbeck process with a time constant of 10 seconds; this lead to realistic trajectories with changes of direction, see *methods* for details. (We will use the metaphor of a virtual animal for conceptual ease, although we really mean that we moved the bump-like external input to our model neurons). Every three minutes of simulation time we stopped the animal at its current location for three seconds and removed the place field input. This led to spontaneous bursting via a synaptic depression-dependent mechanism^27^, reminiscent of sharp-waves seen during awake quiescence and sleep, see Fig.2. Neither the bursting, nor the learning process itself depended qualitatively on the exact amount of time the virtual animal spent moving versus staying still. Synaptic plasticity was allowed to occur during both theta activity and sharp-waves, i.e. it was never “turned-off”.

**Figure 2.**
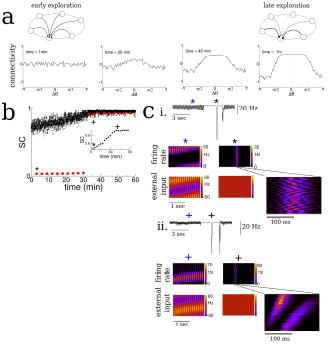
Spatial exploration gives rise to plasticity-dependent emergence of spatio-temporal structure in spontaneous activity. (a) Snapshots of the connectivity during the exploration of a novel environment. During early exploration, the connectivity is not correlated with the place-field ordering, while during late exploration cells with neighboring place-fields are more strongly connected. (b) The mean cross-correlation of the activity of place cells with adjacent place fields on a novel track during exploration (sequential correlation SC). Black: SC during active exploration (theta-activity), Red: SC during spontaneous bursts. Note the sharp transition in the SC of burst activity. Inset: The SC of the total activity binned into 3-minute intervals shows a steady increase preceding the transition. (c) Burst activity exhibits replay after a critical period. i. Early exploration. Top: average input to place cells before (blue star), during (black star) and after period of “quiet wakefulness”. Bottom: space-time plots of the place cell input and firing rate. Note the disordered spatio-temporal structure of the burst activity. ii. Later exploration. After a critical transition time bursts exhibit sequential replay of activity from the novel track. Note the sequential structure of the burst activity.

As the virtual animal explored the track, the sequential activation of the place cells led to changes in the recurrent connectivity via the plasticity rule, Fig.2a. Whereas the connectivity between cells was initially unstructured on the novel track, it evolved over the span of minutes to tens of minutes such that cells with nearby place fields were more strongly connected than ones with disparate place fields. Furthermore the resulting connectivity was slightly asymmetric, reflecting the bias of the animal’s motion, see Fig.S3. Although the changes we observed in the recurrent connectivity, see Fig.2a, and concomitant changes in place cell activity were continuous and smooth during the duration of the simulation, there was a dramatic and sharp transition in the structure of the burst activity during awake quiescence. We quantified this transition by measuring the mean pairwise cross-correlation of cells with neighboring place fields *on the novel track*. Such cells are “neighbors” only on the novel track and not on any other track by virtue of the global remapping or randomization of place-fields. We call this measure the sequential correlation (SC) because it is high when there is a properly ordered sequence of place-cell activations on any given track. The SC during theta-activity (exploration) was already non-zero at the outset of the simulation by virtue of the external place-field input which generates sequential activity, black squares Fig.2b. On the other hand, the SC was initially near zero during spontaneous bursts, red circles Fig.2b. Interestingly the SC abruptly increased during burst activity at a critical timeband remained elevated for the duration of the simulation. Note that the SC for theta-activity also showed a steady increase leading up to this transition, as did the SC when the total activity was taken into account, without discerning between epochs of movement or awake quiescence, see Fig.2b inset. Space-time plots of place-cell activity show that the abrupt increase in SC coincides with a transition in the spatio-temporal structure of the bursts: it is initially uncorrelated with the ordering of cells on the novel track, Fig.2c i., and after the transition there is a clear sequential activation reminiscent of so-called “replay” activity, Fig.2c ii. In fact, before the transition the bursts are highly structured, but this structure is correlated with a previously explored track and not the novel one. If the learning process was carried on for much longer times all relevant network properties, such as connectivity and mean firing rates, saturated, see Fig.S4.

### The transition in the structure of bursts is strongly dependent on the shape of the plasticity rule and the frequency of “theta” modulation

We hypothesized that changes in the recurrent connections between place cells in our model during exploration were shaping the spontaneous activity and driving the transition to replay-like activity. The connectivity profile at any point in time could be decomposed into a series of spatial Fourier modes, Fig.3a. We tracked these modes in time and discovered that the transition in burst activity always occurred when the coefficient of the first even mode reached a critical value. Specifically, we conducted additional simulations in which we increased the firing rate of the feedforward inputs which drove place cell activity which in turn led the transition in burst activity to occur at earlier times, see shaded regions in Fig.3b. In fact, a theoretical analysis of our model shows that there is an instability of the spontaneous activity to bursts when the even mode of the connectivity times the gain in the neuronal transfer function is greater than a critical value, see methods and *supplementary information (SI)*. In our simulations the gain in the transfer function was nearly always the same during bursts because it depended on the mean input to the network during quiescence, which was constant. Therefore it was principally the even mode of the recurrent connectivity which determined the time of the transition. For this reason, the transition to replay occurred when the coefficient of the even mode reached a critical value as seen in Fig.3c. Note that there was no such dependence on the odd mode (dotted lines in Fig.3c).

**Figure 3.**
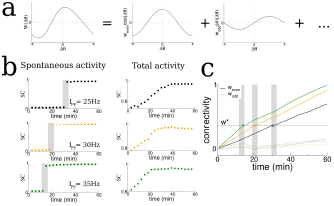
The transition in the structure of burst activity occurs when the even component of the recurrent connectivity reaches a critical value. This mode grows only for STDP rules with dominant potentiation, and grows fastest when periodic modulation is in the theta range. (a) The profile of the recurrent connectivity at any point in time can be decomposed into a series of even (cosine) and odd (sine) modes. Only the first two modes are shown. (b) As the amplitude of the place-field input *I* _*PF*_ is increased from 25Hz (black), 30Hz (orange) and 35Hz (green) the transition shifts to earlier times. The SC of the SWR activity shows a sharp transition (shaded grey bars), while the total activity displays a smooth increase leading up to the transition. (c) In all cases the value of the even mode reaches approximately the same critical value at the time of the transition (shaded grey bars). The growth of the odd mode (dotted lines) is much more irregular, and its value does not correlate with the transition time.

We then asked how the growth of the even mode depended on the details of the plasticity rule and the frequency of periodic modulation of the place cell activity. We found that the growth of the even mode depended strongly on the frequency, peaking in the theta range and decreasing by orders of magnitude at low and high frequencies, Fig.4i. This occurred independent of the details of the animal’s motion, e.g. compare Fig.4i and FigS6e for irregular versus purely clockwise motion respectively. On the other hand the odd mode, which represents a directional bias in the recurrent connectivity, closely tracks the cumulative bias in the animal’s motion, see Fig.S3, and is independent of the forcing frequency to leading order, see methods and *SI* for details. The strong dependence of the growth of the even mode on frequency, meant that, for simulations of up to 1hr, transitions in the burst activity were observed only when the frequency was in the theta range, Fig.4aii. Furthermore, the even mode only grew, and transitions only occurred, when the plasticity rule had dominant potentiation at short latencies. For a perfectly anti-symmetric plasticity rule, and for a rule with dominant inhibition at short latencies, the even mode did not change and even decreased, respectively, Figs.4b and 4c. In the latter case bursts were suppressed entirely (not shown).

**Figure 4.**
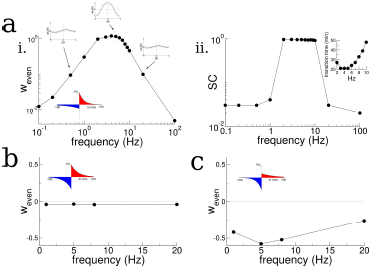
Theta-modulation accelerates emergence of connectivity mode which drives replay. (a) A temporal asymmetric plasticity rule with dominant potentiation at short latencies. i. The amplitude of the even mode of the connectivity after 1hr of exploration, as a function of the modulation frequency. The even mode grows maximally over a range of approximately 1-10Hz and it is strongly attenuated at lower and higher frequencies. Note the logarithmic scale. On the other hand, the growth of the odd mode is largely independent of the modulation frequency, see text for details. ii. The degree of SC of spontaneous bursting after 1hr of exploration. Replay occurs only when the frequency lies between 2-8Hz. Inset: The time at which a transition in the SC takes place as a function of frequency, for times up to 1hr. (b) A purely anti-symmetric plasticity rule leads to recurrent connectivity which has only odd Fourier modes (red squares). There is no increase or transition in the SC of bursts in these simulations even after 1hr. (c) An asymmetric plasticity rule with depression dominating at short latencies leads to connectivity with a negative amplitude of the even mode, i.e. recurrent excitation is weaker between pairs of place cells with overlapping place fields than those with widely separated place fields. In this case bursts are actually completely suppressed (not shown).

### A self-consistent theory to explain how the interplay between theta-modulation and the plasticity rule govern changes in recurrent connectivity

We sought to understand the mechanism underlying the evolution of the recurrent connectivity seen in simulations by studying a simplified, linear version of our network, which was amenable to analysis. In addition, we made use of the fact that the step-wise increases in synaptic strength due to the plasticity rule can be approximated as a smooth process as long as plasticity occurs much more slowly than the firing rate dynamics. When this is the case, the rate of change of the synaptic weight from place cell with place field centered at *θ’* to one with place field at *θ* can be written as

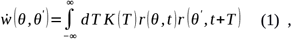

where K(T) is the change in the synaptic weight according to the plasticity rule given a spike pair with latency T^22^. This equation reflects the fact that the total change in the synaptic weight is the sum of all the pairwise contributions from the pre- and post-synaptic cells, with each pair of spikes weighted by the plasticity rule with the appropriate latency. The firing rates are then found self-consistently from the firing rate equation, Eq.5. In the end we find that the evolution of the recurrent connectivity can be approximated by taking

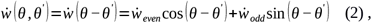

where the growth rates of the even and odd modes 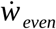 and 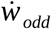 are functions of the STDP rule parameters, the velocity of the animal and the frequency of periodic modulation, see *STAR methods* for details. It turns out it is possible to understand these dependencies intuitively and comprehensively without having to study the analytical formulas. Specifically, if we wish to isolate the growth rate of the even mode, which is responsible for driving the emergence of replay in the burst, we can consider place cell pairs where *θ*=*θ’*, i.e. pairs with overlapping place fields. When this is the case we can combine Eqs.(1) and (2) to yield

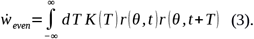

By appropriately averaging both sides of the equation in time (see methods) the product of the rates in the integral becomes the autocorrelation (AC) of place-cell activity, and so we can write

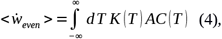

where the brackets indicate averaging over the fast time-scale dynamics.

Therefore the growth rate of the even mode is found by multiplying the AC of place-cell activity by the window for plasticity, and integrating. Because the effect of periodic modulation on the AC is straightforward, we can determine graphically how the frequency of modulation interacts with the plasticity rule to drive changes in the burst structure.

We first note that if there is no periodic modulation of place-cell activity then the AC will simply reflect the movement of the animal. This will lead to a very broad AC compared to the time-scale of plasticity. For example, if we assume that the width of the place field is a fraction of the track length (as in our model), then a rat running between 5 and 50cm/sec on a multi-meter track would have an AC which decays on the order of between seconds and tens of seconds. Therefore, the AC is essentially constant compared to a typical plasticity window of between tens and hundreds of milliseconds, and the integral in Eq.(4) will give nearly zero. Periodically modulating place-cell activity will increase the growth rate as long as potentiation is dominant at short latencies, Fig.5a. Slow modulation will bias the integral in Eq.(4) toward the potentiation lobe of the STDP (Fig5a left and middle, top), while in an optimal range of frequencies the peaks and troughs of the AC will maximally capture potentiation *and* flip the sign of the depression lobe (Fig5a left and middle, middle). Finally, at higher frequencies the plasticity rule undergoes multiple sign flips on a fast time scale, which again gives a near zero value for Eq.(4) (Fig.5a, left and middle, bottom). This means that the maximal growth rate of the even mode occurs for an intermediate range of frequencies: those which modulate place-cell activity on a time scale commensurate with the window for plasticity, Fig5a right. Note that this has nothing to do with the overall rate of plasticity, which is the same independent of the modulation frequency. That is, even if the AC is flat, recurrent synapses undergo large numbers of potentiations and depressions. Rather, periodic modulation serves to organize the structure of synaptic plasticity at the network level by preferentially strengthening connections between place-cells with overlapping or nearby place field and weakening others. The optimal range of frequencies for growth of the even mode depends only weakly on the ratio of the width of the potentiation to that of the depression lobe, Fig.5c, but significantly on the total width, Fig.5d. Allowing for triplet interactions (as opposed to just pairwise) in the plasticity rule increases the overall growth rate but does not alter the range of optimal frequencies, Fig.5e. On the other hand, the theory predicts that the growth rate of the odd mode is only weakly dependent on the modulation frequency (Fig.5b) as is seen in simulations (Fig.S6e), and can be understood by considering Eq.(2) with *θ-θ’* =*π* /2. In this case the growth rate depends on the product of the plasticity rule with the cross-correlation (CC) of cells with disparate place fields. When there is an overall direction bias in motion then the CC will have peak shifted from zero-lag and the product with the STDP rule reliably gives a positive (or negative) growth rate. When there is very little motion bias the CC will be nearly flat, yielding little growth in the odd mode and the resulting connectivity will be highly symmetric.

**Figure 5.**
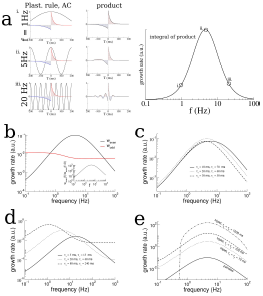
Analysis of a linear firing rate model with plasticity reveals the mechanism by which the plasticity rule and periodic modulation shape connectivity. (a) The growth rate of the symmetric mode is proportional to the integral of the product of the autocorrelation (AC) of place cell activity with the plasticity window (PW). Left: The AC of place cell activity overlaid on the PW with dominant potentiation at short latencies. Middle: Product of the PW and the AC. Right: The growth rate of the symmetric (cosine) mode of the connectivity as a function of frequency. i. When the activity is modulated at 1Hz (top), the product returns nearly the original plasticity window, which being balanced yields near-zero growth rate. ii. For 5Hz the potentiating lobe is maintained and some of the depression lobe changes sign and becomes potentiating leading to higher growth rate. iii. For 20Hz the plasticity window undergoes sign reversals at a rate faster than the width of the lobes, meaning the integral is again near zero. (b) The growth rate of the even (cosine) and odd (sine) spatial Fourier coefficients as a function of the modulation frequency (black curve is the same as in (a) except on a log-log scale). Inset: The growth rate of the even mode normalized by its value for no periodic modulation. Rule parameters are *A*_+_=0.1, *τ*_+_=20 *ms, A*_-_=0.1/3, *τ*_-_=60 *ms.* (c) Increased growth rate in the theta range does not require fine tuning. (d) The frequency at which growth is maximal depends on the overall width of the plasticity window. Broader windows favor slower frequencies. (e) A triplet rule increases the growth rate at all frequencies compared to the pairwise rule, but does not significantly shift the optimal frequency range. The parameters for pairwise interactions are as before. The time constants for triplet interactions are indicated on the figure, while the remaining parameters are chosen to make the rule balanced.

Finally, it is also clear from Eq.4 that a perfectly anti-symmetric plasticity rule will lead to no growth in the even mode as the product of the rule and the AC will itself be an odd function, see Fig4b. A rule with dominant depression at short latencies will similarly yield a growth rate which is precisely the inverse of that shown in Fig.5a. This causes a decay in the even mode resulting in a connectivity pattern with locally dominant inhibition, see Fig.4c.

### Sparse coding allows for the replay of activity from multiple explored tracks during bursts

The spatio-temporal structure of sharp-wave-like bursts in our model reflected the sequential ordering of place fields on the most recently explored track; correlation with earlier tracks was erased or greatly reduced. In reality, only a fraction of place cells have well-defined place fields in any given environment, i.e. coding is sparse^6,28^. We incorporated this sparse-coding strategy to our model by providing place-field input to only one half of the total neurons in the network; the other half received constant input. These place cells were chosen randomly from one track to the next and place field locations were assigned randomly as before. Therefore the overlap in the population of place-cells between any two tracks was also fifty percent. We then allowed the network to evolve by having the virtual animal explore thirty distinct tracks, each for 1hr of simulation time, as before. The resulting matrix of synaptic connections was correlated with the ordering of place-cells in several past explored environments, Fig.6a. The amplitude of the even mode, which is responsible for generating spontaneous replay, decayed roughly exponentially as a function of the recency of the explored track, Fig.6a inset. This suggested that the replay dynamics in the network with sparse coding might be correlated with several previously explored track simultaneously^27^. Indeed, when the network was driven with a global, non-selective input, the replay was strongly correlated with the past two explored environments, Fig.6b, whereas the correlation with earlier environments was negligible. On the other hand, when the constant external drive was selective to the subset of place-cells on any given track, replay activity was robustly correlated with activity on that track only, Fig.6c. Such selective input may originate in cortical circuits which store environment-specific sensory information as stable patterns of activity; these patterns correspond to attracting fixed points in network models of long-term memory storage and memory recall^29–31^. Spontaneous switching between cortical representations during slow-wave activity would therefore engage distinct hippocampal replay patterns, which in turn could strengthen and consolidate the corresponding cortical memory trace.

**Figure 6.**
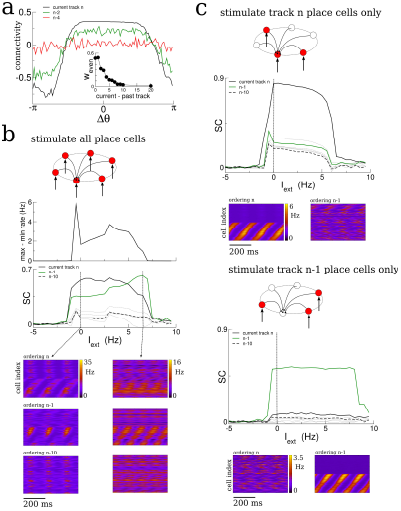
Sparse coding allows for the replay of multiple previously experienced environments. (a) The connectivity profile after exploration of 30 environments in a network in which one half of the neurons are place cells on any given track. The same connectivity can be visualized using the ordering of the place cells on the most recently explored track (track n, black curve) or using the ordering of past explored tracks (green and red curves). The spatial structure which emerges during exploration on any given track eventually gets overwritten as new tracks are explored. Nonetheless the connectivity stores structure from several past explored tracks simultaneously. Inset: The amplitude of the even Fourier mode as a function of the recency of the corresponding track. (b) A global stimulus applied to all the neurons in the network generates replay which is highly correlated with the past two tracks. (c) Selective stimulation of only those neurons which were place cells in the most recently explored track (top) or next-to-last track (bottom) generates replay which is highly correlated only with the corresponding environment. N = 200 neurons in all simulations.

Therefore, sparse coding causes the synaptic connectivity matrix to be simultaneously correlated with place field orderings from multiple tracks and allows for robust replay of activity from those tracks given appropriate inputs. The number of different environments which could be simultaneously encoded in replay depended on model details, particularly the coding level, i.e. the fraction of active place cells in any given environment (not shown)^32^.

### Experimental evidence for a transition to replay

Pairwise reactivations of CA1 place-cells with overlapping place fields during awake SWRs improve during exploration; they are stronger during late exploration than early exploration^33^. This holds true not only for pairwise correlations, but also reactivations of entire neuronal ensembles, at least on linear tracks^34^. More recent work has shown that significant replay events during awake SWRs in rats running along a three-arm maze emerge abruptly only after a certain number of runs^12^. These results are consistent with our model predictions. We additionally sought to directly test for a time-resolved increase in SC in neuronal data. We looked for this increase in multi-unit recordings of place-cell activity from the hippocampus of rats exploring novel, periodic tracks^19^. We first identified cells with well-defined place fields by extracting the coefficients and phase of the first two spatial Fourier modes of their time-averaged activity as a function of the normalized distance along the track (in radians), see Fig.S7, S8 and S10. We kept only those cells for which the ratio of the coefficients, the Tuning Index (TI), exceeded the threshold of one, indicating strong spatially selective activity, see *methods* for details. We then ordered the cells according to their phase (approximately the position of peak firing). The SC of activity over the total duration of the experiment using this ordering was significantly higher than 5,000 randomly reshuffled orderings which on average gave SC = 0, see Fig.7a and b. When the medial septum (MS) was inactivated via muscimol the SC did not exhibit any dynamics as a function of time, Fig.7c. However, once the animal recovered from the muscimol the SC using the proper phase ordering exhibited an initial ramp over the first ten minutes of exploration and then remained high (significant difference between first point and all others, t-test with multiple-comparison Bonferroni correction, p-values < 0.004 and between second and third points), see Fig.7d, solid circles. This is consistent with the model result which showed a similar ramping increase when the total activity (and not just bursts) was considered, see e.g. Fig.2c inset. On the other hand, the SC computed for shuffled phases showed no dynamics and remained close to zero, see Fig.7c,d, open squares. This indicates that there are no global changes in neuronal correlations during exploration, which could occur, for example, due to slow changes in theta modulation or neuronal excitability. Rather, there is a sustained increase in pairwise correlations only between those neurons which encode nearby places, and only when strong theta modulation is present. This finding also held when we considered the more lax criterion *TI* ⩾1/2, see Fig.S9 (rat A991). In a second animal there was an insufficient number of well-tuned units to repeat the analysis, see Fig.S9 (rat A992). One possible confound in attributing this increase in correlation to theta modulation alone, is the fact that the firing rates of place-cells during MS inactivation were lower on average than during the post-muscimol experiment. Lower firing rates in the computational model lead to lower rates of plasticity. However, this difference in firing rates was not significant for the well-tuned neurons shown in Fig.6 (*TI ≥* 1), (t-test, p = 0.30). An initial, sustained increase in SC was also observed in data from a separate experiment in which an animal was first exposed to a novel hexagonal track, Fig.6e (difference between first point and all others, t-test with correction for multiple comparison, p-values < 10^−6^). Furthermore, no such increase was found on subsequent sessions on the same track, indicating that the change in correlation only occurred when the environment was novel. This result held when we considered the more lax criterion *TI* ⩾1/2 and also when all units were used, regardless of tuning, see Fig.S10a (rat A992). In a second rat there were too few well-tuned cells to use the criterion *TI ≥* 1, but an initial and sustained increase in SC was seen for *TI* ⩾1/2 and also using all units, although this correlation was not always significantly different from that calculated from shuffled orderings of the cells (unshaded points in Fig.S10a, rat A991).

**Figure 7.**
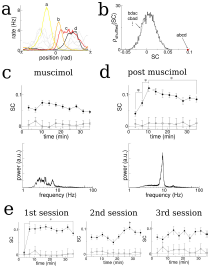
The SC of place cells in rat hippocampus during the exploration of a novel track shows an initial increase and plateau, but only when theta is present. (a) Sample firing rate profiles from place cells recorded from CA1 of rat hippocampus during exploration of a novel periodic track with an illustration of how cells can be ordered according to their phases. (b) The SC calculated over the entirety of the experiment (35 min) given the correct ordering (red dot) and for 5000 reshuffled orderings. (c) The time course of SC given the proper ordering (solid symbols) does not show any dynamics when the medial septum is reversibly inactivated with muscimol. Note, however, the clear separation with the shuffled data (open symbols), indicating that place fields are intact even though theta is disrupted. (d) After a rest period the animal is placed back onto the same track; the SC now exhibits a significant increase given the proper ordering (solid symbols) over the first 10 minutes of exploration and then plateaus. Note that the bottom plots in (c) and (d) show the power spectrum of hippocampal LFP (CA1), indicating a large reduction of theta power in the muscimol condition. (e) When the animal is allowed to explore a novel track and then placed back on the track for a second and third session, there is a significant increase in the SC only during the first session. Error bars are S.E.M.

## Discussion

### Summary

We have presented a computational model of a hippocampal place-cell network in order to investigate how the exploration of novel environments shapes the patterns of recurrent synaptic connectivity. Because place-fields remap randomly from one environment to the next, the recurrent connectivity, shaped by previous learning, is initially uncorrelated with place-cell activity in a novel environment. Our major finding is that the rate at which spatial correlations in the connectivity emerge during exploration depends almost entirely on the product of the autocorrelation of place-cell activity and the window for plasticity. The integral of this product determines the growth rate of a global, network-wide pattern of synaptic connectivity, which results in strong local recurrence and long-range competition. It is this mode which drives spontaneous replay activity in our model network. The growth rate of this mode is maximum when place-cell activity is periodically modulated on a time-scale commensurate with the plasticity rule, which for realistic time constants yields frequencies in the theta range. Furthermore, lower and higher frequencies than theta lead to learning rates which are orders of magnitude slower. This suggests that the role of theta is to accelerate learning. Note that the overall rate of plasticity is not affected by the presence of oscillations. The number of spike pairs, and hence the number of potentiations and depressions, depends only on the firing rates. Rather, theta oscillations generate repeated pre-post pairings in both directions, which coupled with a plasticity rule with dominant potentiation at short latencies bias plasticity toward potentiating events for neurons with neighboring place fields. One signature of this mechanism is a continuous increase in the pairwise cross-correlation in the activity of neighboring place-cells leading up to a critical time. We have found evidence consistent with this by analyzing the activity of simultaneously recorded hippocampal place cells in a rat during the exploration of a novel track.

### The assumption of plasticity at recurrent synapses

In our model we have assumed that plasticity occurs only in the recurrent excitatory synaptic connections, and not in the feed-forward inputs. Therefore we also assume that the place-field input, which peaks at the spatial position of the virtual animal at given moment in time, is itself stable. In fact, consistent with this assumption, it seems most place cells are active from the outset of exploration of a new environment, although see^35–37^. Furthermore cells tend to exhibit only subtle changes in the size and shape of their place fields over time^38,39^, also consistent with our model, see Fig.S6b. On the other hand, it has been shown in area CA1 that some place-cells exhibit place-fields only after several minutes of exploration^36,40^. Recent intracellular recordings indicate that appearance of these “hidden” place-fields requires the coincidence of active dendritic events and synaptic input via the Schaeffer collaterals^41–43^. It may be that this mechanism for place-cell “generation” is particularly salient in cells of area CA1 by virtue of being uniquely positioned to compare and integrate both entorhinal and hippocampal inputs. In any case the strongly recurrent nature of the network we study may make it a more relevant model for circuits in area CA3.

Nonetheless we would expect that changes in spiking activity arising in CA3 due to plasticity in the recurrent connectivity, as in our model, would be reflected in similar changes in the spiking activity of CA1 cells due to the direct inputs via the Schaeffer collaterals. More specifically, in contrast to plasticity mechanisms leading to the formation of place cells themselves, here we have modeled on how plasticity shapes the recurrent connections between already-formed place cells. We find that pairwise correlations between place cells with nearby preferred locations increases during exploration of a novel environment. Assuming such an increase occurs in a strongly recurrent circuit in CA3, we would also expect to observe an increase in correlation in target CA1 pyramidal cells, as long as there exists some mapping from CA3 place cells to CA1 place cells. Such a mapping could be a simple random projection or a more ordered relationship. Recent work suggests that place fields of CA1 pyramidal cells on linear tracks are built up of a weighted sum of CA3 inputs from positions surrounding the relevant place field^43^; in such a case the increased correlation in CA3 activity would be shared by CA1 output. Indeed, the data we have analyzed from place cells of area CA1 show increase sequential correlation as predicted by our recurrent model. A recent computational model of STDP-induced formation of place fields in CA1 cells via the feed-forward excitatory connections from CA3 (Schaeffer Collaterals) suggests a role for theta in speeding up place cell formation^44^. This raises the intriguing suggestion that theta may play a key role both in place-cell formation through plasticity of feed-forward inputs, and also in the emergence of replay through plasticity of recurrent synaptic connections as indicated by our work here.

### Remapping of place fields for different directions of motion on the same track

Hippocampal place cells actually exhibit global remapping of their place fields depending on the direction of motion of the animal on linear tracks^45,46^. This is perhaps not surprising given that the behaviorally relevant information for the animal is not just the position along the track but also the way in which it is facing; e.g. this determines how far away a potential reward is, if located at one or both ends of the track. Studies using periodic tracks have shown no such global remapping, but rather some degree of rate remapping^47^, i.e. the direction of motion affects the shape and amplitude of place fields, but not their position. In the data we have analyzed there is very weak remapping, see Figs.S7c and Fig.S8b, and so the assumption of invariance of place field to direction of motion is a good one. In our model we have made this assumption. The consequence of this is that while exclusively clockwise (CW) or counter-clockwise (CCW) motion will lead to highly asymmetric recurrent connectivity Fig.S6b, exploration of both directions will give rise to much more symmetric connectivity, see Fig.2a. In practice any trajectory over a finite amount of time will have a directional bias; in the data we have analyzed the rat spends 54% of the time moving CW and 46% CCW, and this will necessarily lead to asymmetries in the connectivity. In linear tracks, due to the global remapping such asymmetries should be even more pronounced.

### Forward versus backward replay

The inevitable asymmetry in the recurrent connectivity of our model place-cell network due to plasticity during exploration strongly biases spontaneous activity. On a periodic track this replay would be exclusively CW or CCW depending on the corresponding bias in motion during exploration, while learning on a linear track would always produce forward replay. Previous work has shown that perfectly symmetric connectivity can give rise to both forward and backward replay in equal measures, due to spontaneous symmetry breaking of activity^27^. We would argue that such symmetric connectivity is not robust for the reasons given above, although we cannot rule out the existence of homeostatic mechanisms which would conspire to make it so. Rather, we propose here an alternative mechanism for generating backward replay given asymmetric connectivity based on local sensory input. Specifically, if global input to our model network is constant then replay occurs only in one direction. However, if a localized bump of input is provided to the network, synapses to downstream neurons (in the sense of the asymmetric bias of the connectivity) become rapidly depressed. This prevents the spontaneous activity from propagating forward and forces it to propagate backward, see Fig.S5. In fact, in experiment, when local spatial input is absent, e.g. when the animal is sleeping in a rest box, forward replay is predominant^48^. On the other hand, both backward and forward replay are observed when the animal is awake but quiescent on a given track. This is precisely when locally sensory cues are available to the animal, and could potentially shape spontaneous replay events. In fact recent work shows that some neurons in area CA2 fire more strongly during awake quiescence than during exploration^49^; they may be providing information regarding local sensory cues. In our scenario the mechanisms leading to forward versus backward replay are distinct and therefore in principle relevant replay statistics such as replay velocity and spiking intensity should also be different.

### Robustness to changes in the plasticity model and to the presence of spike correlations

Here we have considered a simple phenomenological model of plasticity which depends on the timing of spike pairs. Taking into account spike triplets as opposed to only pairs does not alter our findings qualitatively, see Fig.5e, although we are necessarily ensuring a balanced rule through our choice of parameters. Fits of heuristic spike timing-dependent models to data from slice experiments yield parameter values which do not lead to balanced rules such as the ones we have used here^24^. In a recurrent network model, unbalanced plasticity rules would quickly lead to synapses either saturating to maximum efficacy or vanishing completely; non-trivial network structure is therefore not possible with such rules. One solution to this problem is to complement the heuristic, unbalanced Hebbian rule, extracted from slice experiments, with a non-Hebbian heterosynaptic rule, as well as additional, slower homeostatic mechanisms^26^. However, heuristic rules describing synaptic plasticity *in-vivo* are still largely unknown. Therefore one reasonable approach to studying plasticity in recurrent networks *in-vivo* is to choose the simplest possible rule which allows for the emergence of non-trivial structure; this has been our approach here. It remains to be studied how more realistic voltage- or calcium-based plasticity rules interact with the theta-modulation to affect learning in recurrent networks^50,51^, although at a single-synapse one can find qualitatively similar regimes for an array of plasticity rules in the presence of pre- and post-synaptic oscillations^52^.

Our results clearly do not depend on the actual spike timing since our model neurons generate spikes as Poisson processes; rather, all lasting changes in the connectivity are due to time-varying modulations of the firing rates. In fact, recent work with a spiking neuron model suggests that such modulations in the firing rate, as opposed to exact spike timing, are sufficient to explain the effect of plasticity from STDP and more realistic calcium-based plasticity rules in general^25^. In any case the contribution of pairwise spike correlations to the evolution of the recurrent connectivity can be formally taken into account in Eq.4, i.e. via its affect on the auto-correlation of place-cell activity.

### Other network models of place-cell activity

Recurrent network models for place-cell activity provide a parsimonious explanation for electrophysiological phenomena associated with exploratory behavior as well as the generation of sharp-wave bursts during awake quiescence and sleep^27, 53–56^. In recent theoretical work^27^ sharp-wave-like bursts were generated spontaneously by virtue of the spatial modulation of the recurrent connectivity, which drives an instability to traveling waves in the absence of place-field input. The presence of short-term depression modulates the amplitude of the waves, leading to narrow bursts. This is the same mechanism we have used here. Alternatively, recent work with a biophysically detailed spiking network model focused on the role of nonlinear dendritic integration on the generation of replay during SWRs^56^. In that work the authors found that the exploration of a virtual linear track in the presence of pairwise STDP lead to highly asymmetric connectivity; this could generate replay activity given a sufficiently synchronous external input which recruited nonlinear dendritic events. In our work, we have sought to explain the replay as an emergent phenomenon which depends only on the network-wide organization of synaptic structure. In doing so we have considered a simple stochastic firing rate model which allowed us to fully characterize how interplay between the plasticity rule and the place-cell activity affects learning. Nonetheless, a detailed reproduction of the phenomenology of SWRs certainly requires mechanisms we have not included here. In particular, while our model generates sharp-wave-like bursts, it does not generate high-frequency ripples, which are likely generated by networks of inhibitory interneurons in CA1^57^.

### Spatial learning

It seems reasonable that the learning of tasks which depend on spatial information require the formation of an internal representation of the relevant environment. This is the process we have studied here. While we have not modeled any particular cognitive task, we propose that the network-wide organization of synaptic structure, in order that it be in concordance with the place-field distribution of place cells, should be a necessary step in spatial learning tasks. Our results suggest that this process is dramatically sped up by modulating place-cell activity in the theta range, which is one possible role of this prominent rhythm. More generally, for learning to occur on behaviorally relevant time scales, neuronal activity must vary on a time-scale commensurate with synaptic plasticity mechanisms. One means of achieving this is to have a broad window for synaptic plasticity, on the order of seconds, so-called behavioral time-scale plasticity (Bittner et al. 2017). Alternatively, internally generated rhythms may serve to modulate neuronal activity on non-behavioral time-scales, with a similar effect.

## Author Contributions

P.T. developed theory and ran numerical simulations. B.R. analyzed data. Y.W. provided data and assistance with data analysis. A.R. developed theory, ran simulations, analyzed data and wrote the paper with input from the other authors.

## Acknowledgements

A.R. acknowledges grants from the Spanish Ministry of Economics and Competitiveness, grants no. BFU2012-33413 and MTM2015-71509.

## Methods

### Method Details

We simulated a model of n excitatory neurons with global inhibitory feedback. The firing rate of a neuron i evolved according to 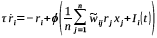 where 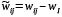 is the effective connectivity from a cell j to a cell i and consists of a recurrent excitatory synaptic weight and a global inhibitory feedback. We take the neuronal transfer function to be *ϕ*(*I*)=*α* ln (1+ *e*^*I* /*α*^) where *α* =1 *Hz* for all simulations^27^. Therefore, both the input and output of the function have units of firing rate. The synaptic depression variable *x*_*i*_ obeys 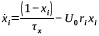. The external input is *I*_*i*_ (*t*)=*I*_0_ + *I*_*PF*_ cos (*θ*_*i*_ - *θ* (*t*)) (1+*I*_*theta*_ cos (2 *π ft*)) where *θ*_*i*_ is the place field position of neuron i in radians, *θ* (*t*) is the position of the “virtual animal”, *I* _*PF*_ is the amplitude of the place-field input. To model theta-modulation we multiplied the place field input by a periodic signal with frequency *f* and amplitude *I*_*theta*_. This type of multiplicative modulation is seen in intracellular recordings in-vivo, e.g. see Figs.1 and 5 in (Harvey et al. 2009) ^3^ and Fig.4 in (Lee et al. 2012)^41^. A cell i generates a spike in a time interval dt with probability *r*_*i*_ *dt*. Plasticity occurs for every spike pair between cells i and j:

*w*_*ij*_ *→w*_*ij*_+ *Δw*_*ij*_ where *Δ w*_*ij*_= *A*_+_ exp (*-T* /*τ*_+_) if *T* =*t*_*i*_ *-t* _*j*_ >0 else *Δ w*_*ij*_=*- A*_-_ exp (*T* /*τ*_-_).

We implement this as in (Pfister and Gerstner 2006)^24^. We furthermore set a minimum value for synapses at zero and a maximum value *w*_*max*_. We model distinct tracks by spacing place fields uniformly and assigning them randomly to cells. Therefore the ordering of *θ*_*i*_ s from track to track are random and uncorrelated. In all simulations the number of neurons is n = 100, except for those in Fig.6 for which n = 200. Note that given the stochastic nature of the simulations and the relatively small system size, different realizations will give rise to quantitatively different results, e.g. jitter in the transition time, variability in the shape of the SC. Nonetheless, the curves shown are typical and qualitatively representative.

### Model analysis

#### How theta modulation and STDP shape recurrent connectivity

We consider a continuum limit of the network neglecting synaptic depression, in which the firing

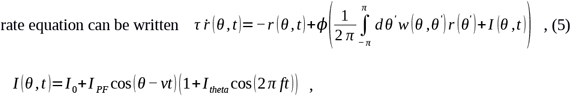

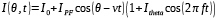

and assuming the plasticity is slow compared to the rate dynamics we can write

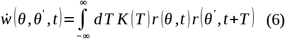

If we take the transfer function ϕ to be linear we can solve the equations self-consistently for the rates and the weights where 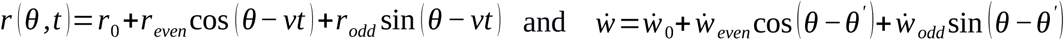. Note that this solution is strictly only valid when the synaptic weight changes *A*_+_ and *A*_-_ are small. Formally, we define a small parameter, proportional to the synaptic weight change, as well as a slow-time, thereby separating the fast (neuronal) and slow (connectivity) time scales. This allows us to assume the connectivity is constant on a fast time scale, in which case Eq.5 is a simple linear system, see SI for details. Finally, the slow evolution in Eq.6 is driven solely by the terms arising from the product of the rates which have non-zero time average. Other terms are eliminated by averaging over the fast time scale, again see SI for a detailed derivation.

We choose a balanced STDP rule to avoid unbounded growth or decay to zero in the mean weights, i.e. *A*_+_ *τ*_+_= *A*_-_ *τ*_-_. Then 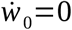,

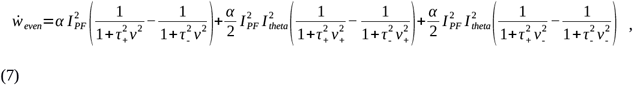

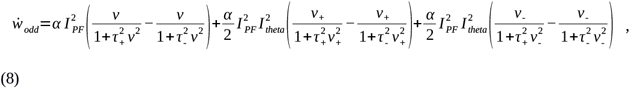

where *v+* =*v*+ 2 *π f* and *v* =*v-* 2 *π f* and 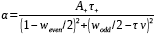. For modulation frequencies above 1Hz we have 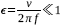 and the growth rates simplify to give

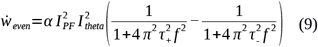

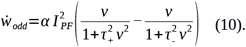

This shows explicitly that the growth of the even mode is strongly dependent on the amplitude and frequency f of periodic modulation, while that of the odd mode is independent of these to leading order. See SI for a detailed derivation. Interestingly, in a single-synapse model of STDP, maximal potentiation is achieved when the pre- and post-synaptic firing rates are modulated at a frequency commensurate with the plasticity window^52^. However, the growth rate of the synapse depends not on a difference of terms, as in Eq.8, but rather on a product. Therefore, in that work, there is no sensitivity to the exact shape of the STDP window, i.e. dominant potentiation or dominant depression at short latencies, or perfectly anti-symmetric.

### The emergence of replay

When the virtual animal is first exposed to a novel track, the recurrent connectivity w is initially unorganized with respect to the ordering of place fields, i.e. *w*_*even*_=*w*_*odd*_=0. We have seen in the last section how exploration and theta-activity cause these modes to grow in time. If we remove the place-field input at different points in time during exploration, i.e. *I* =*I* _0_, we can predict when replay should first emerge. To do this we consider the stability of the homogeneous state in firing rates to spatio-temporal perturbations, i.e. 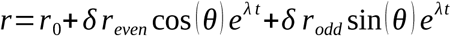, where *δ r* _*even*_ and *δ r* _*odd*_ are small. Plugging this into Eq.(5) yields the characteristic equation for the eigenvalue *λ*, 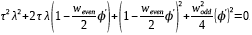. From this we can see that there can be no instabilities to stationary bumps *λ*=0 but that traveling waves can emerge (*λ*=*iω*) when 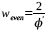 and they have velocity 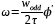. Therefore, exploration will cause the even mode of the connectivity to grow until a critical value is reached, at which point traveling waves correlated with activity on the novel track will emerge spontaneously. This argument still holds when we include the effect of synaptic depression, although the calculation is more involved, see SI.

### Reduction of the plasticity rule to integral of plasticity window times AC of place cell activity

To go from Eq.3 to Eq.4 we must expand the product of the firing rates. For the sake of illustration we will take *r* (*θ, t*)=*r*_0_ +*r*_1_ cos (*θ-vt*) and leave the more general case, with an odd component, for the *SI*. The product

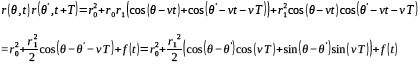

where f(t) is a time-periodic function with period *π* / *v* which could range from 100ms to a few seconds depending on whether v is the velocity of the animal (no modulation) or *v*_+_=*v*+2 *π f*, i.e. theta-modulation. Note that this time-scale is fast compared to the time-scale of plasticity which occurs on the order of minutes. This separation of time-scales allows us to average both sides of Eq.3; if we furthermore set *θ*=*θ*^*’*^ we obtain Eq.4.

### Analysis of data from hippocampal recordings

We analyzed data collected from hippocampal recordings using bilateral silicon probes from two distinct sets of experiments. In the first set two rats had their medial septa reversibly inactivated through injection of muscimol and were then exposed to a novel square, periodic track^19^ for 35 minutes. After a recovery period they were re-introduced onto the same track for an additional 35 minutes. In the second set of experiments two rats were exposed to a novel hexagonal track for 35 minutes in a first session. Thereafter they were placed back onto the same track for two additional 35 minute sessions with a rest period in between each session.

#### Medial septum inactivation experiments

There were a total of n = 124 and n = 128 cells from rats A991 and A992 respectively. In order to identify place cells we first calculated a rate map on the track for each neuron, and then linearized the rate to obtain a one-dimensional place field. We then fit the place field with the function *r*=*r*_0_+ *r*_1_ cos (*θ-ϕ*_1_)+ *r*_2_ cos (2(*θ-ϕ*_2_)) thereby extracting the coefficients of the Fourier modes and their phases, see Fig.S7a and b and S8a. We also extracted the coefficients via Fast-Fourier Transform and found identical results. We did not find any cells for which *r*_2_ was significant compared to the first two coefficients and hence limited our analysis to the first two coefficients and the phase, i.e. the place field was centered at *ϕ*_1_. We excluded all cells for which the tuning index (TI) *r*_1_ / *r*_0_ was below one. We additionally required that the mean firing rate of selected cells over the entire experiment was greater than 0.4Hz. With these criteria we had a total of n = 13 and n = 19 cells for the muscimol and post-muscimol sessions in rat A991 and n = 4 and n = 0 cells for rat A992, see Fig.S9. To ensure we had a sufficient number of neurons we additionally compared the time-resolved SC to the SC from 5,000 shuffled orderings of the cells (t-test, p < 0.005, corrected for multiple comparison). Significant differences indicate that the SC carries spatial specificity as opposed to simply reflecting the degree of global correlation in the network (shaded regions in Fig.S9). The SC for rat A991 was always significantly different from shuffled for all TI, while for rat A992 it was often not so. For this reason we only used the data from rat A991 for these experiments, see Fig.7. We furthermore computed the power spectrum of a simultaneously recorded local field potential (LFP) signal, see Fig.7c and d and Fig.S9.

#### Hexagonal track experiments

There were a total of n = 158 and n = 134 cells from rats A991 and A992 respectively. We selected cells as described above. This left a total of n = 6, 9 and 4 cells and n = 17, 20 and 13 cells for rat A991 and A992 respectively for the three sessions. The SC calculated for rat A991 was not significantly different from that from 5,000 shuffled data sets with random re-orderings of the cells, see non-shaded regions in Fig.S10. Therefore we only considered data from rat A992 for these experiments, see Fig.7e.

### Methods and parameters for figures

*Fig.2* Parameters for simulations are:

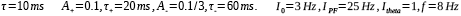

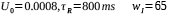

The motion of the animal was modeled by taking the velocity as an Ornstein-Uhlenbeck process. Specifically, the velocity is given by *v* =*v*_0_ +*v*_1_ (*t*) where the mean velocity *v*_0_ =0.5 *rad* / *sec* and 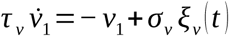 where *ξ*_*v*_ is a Gaussian white-noise process with mean zero and unit variance. The parameters *τ* _*v*_=10 *sec* and *σ* _*v*_=2 *rad* / *sec*. The weight matrix *w*_*ij*_ was trained by simulating exploration on 10 distinct linear tracks for one hour each. The value of *w*_*ij*_ was taken as a constant 40 for all synapses at the beginning of exploration of the first environment and the maximum possible weight was set at *w*_*max*_=80. See Fig.S1 for statistics of the synaptic weights during the training process. For the connectivity profiles, we calculate the mean synaptic weight between pairs of neurons with a difference in place field location *Δθ* at the given time of the snapshot. There are no autapses, but the curve is made continuous at *Δθ*=0 by interpolating between adjacent points. The connectivity is normalized by subtracting the mean and dividing by 40.

To generate the figure in (b) we calculate the SC in 1 sec bins during the entire simulation. Every 180 seconds there is a 3 second period during which the external theta-modulated place field input is removed, in order to model awake quiescence. Activity during this period is spontaneous and is considered to be bursts (black circles). During burst activity we only calculate the SC for the second and third seconds because it takes some time for the place field activity to die away and the burst activity to emerge, e.g. SC for bursts is calculated for seconds 181-182 and 182-183 but not for 180-181. For the simulation in (b) the place field input is simply removed when the animal stops, i.e. the external input is set to a constant value of *I* _0_=3 *Hz* while for (c) it is set to *I* _0_=−0.75 *Hz*. Changing *I* _0_ does not significantly affect the value of SC, only the degree of burstiness of the spontaneous activity, see Fig.S2. The inset in Fig.2b was generated by binning the SC of the total network activity into 3-minute bins. The “LFP” in Fig.2c is the network-averaged input current to the neurons, i.e. the network-average of the argument in the firing rate equations. Fig.3 The curves shown in (a) are a cartoon meant to illustrate how the recurrent connectivity can be decomposed into a spatial Fourier series which include even (cosine) and odd (sine) terms. The amplitude of an even term (its coefficient) can lead to a transition in the network dynamics when it reaches a critical value. In (c) the coefficients *w*_*even*_ and *w*_*odd*_ are the first cosine and sine Fourier coefficients of the mean recurrent connectivity are calculated as in Fig.2 at a given time during the simulation. Parameters in (c) are the same as in Fig.2.

Fig.4 The parameters in (a) and (b) are the same as in Fig.2. Parameters in (c) and (d) are the same as in Fig.2 with the sole exception of the STDP rule. For the anti-symmetric case (c) the parameters are *A*_+_=0.1, *τ*_+_=40 *ms, A*_-_=0.1, *τ*_-_=40 *ms.* while for (d) they are *A*_-_=0.1, *τ*_+_=20 *ms, A*_+_=0.1/ 3, *τ*_-_=60 *ms.*

Fig.5 The virtual animal has explored thirty distinct environments for one hour each. In each environment the coding level is one half, i.e. one half of the neurons are modeled as place cells (randomly assigned place field location from uniform distribution around the track) and the other half receive only constant background input with *I* _0_=0. Plasticity occurs via STDP as before between all cell pairs. Place cells in any given environment are chosen randomly with equal probability; hence the overlap in place-cell representation between any two environments is on average one half. The number of neurons is N = 200, so that in any environment there are still 100 place cells, as in previous simulations.

*Fig.S6* (b) The activity profiles are averages over the five seconds preceding the given time of the snapshot, e.g. for 1 min it is the average activity between 55 and 60 seconds. The average is taken with respect to the peak of the place field input, at *θ*=0.

